# Adaptive introgression in Tibetans spans focal and distributed architectures

**DOI:** 10.64898/2026.09.18.752559

**Authors:** Shen-Ao Liang, Qiong Zhu, Xinyun Gao, Yini Xiao, Kai Liu, Zhengwei Xie, Lian Deng, Xueling Ge, Bing Su, Qiaomei Fu, Yuanting Zheng, Yaoxi He, Shuhua Xu, Lu Chen

## Abstract

The Denisovan-derived *EPAS1* haplotype is a paradigmatic example of adaptive introgression, but whether archaic ancestry contributed beyond such large-effect loci remains unclear. We analyzed genomes from 1,039 Tibetans, Han Chinese, archaic hominins and ancient Tibetan Plateau individuals. Denisovan-derived adaptive variation was highly localized around *EPAS1*, whereas Neanderthal-derived variation was distributed across the genome; temporal and elevational patterns reinforced this contrast. Despite individual-level heterogeneity, Neanderthal-introgressed gene repertoires repeatedly converged on immune, metabolic and signal-transduction pathways. A Tibetan-enriched Neanderthal *PPARGC1A* haplotype was associated with lower fasting glucose and showed concordant support from an independent East Asian type 2 diabetes GWAS. These findings reveal that adaptive introgression can contribute to complex human adaptation through functionally convergent combinations of archaic variants, rather than only through shared large-effect haplotypes.

## Introduction

Genome sequencing has established that present-day human populations retain ancestry from Neanderthals and Denisovans, with the amount and distribution of archaic ancestry varying across populations (*1–14*). A subset of this introgressed variation has been retained by positive selection and contributed to population-specific traits, including immunity, metabolism and environmental adaptation (*15–21*). Many of the best-characterized examples, however, are organized around individual, strongly selected haplotypes, raising a broader question about the architecture of adaptive introgression: must adaptive archaic ancestry be concentrated at shared large-effect loci, or can it instead be distributed across multiple loci that vary among individuals yet converge on common biological functions? Resolving this question is important for understanding how introgressed ancestry contributes to complex adaptation beyond canonical single-locus examples.

Tibetan high-altitude adaptation provides a powerful setting in which to address this question. The Denisovan-derived *EPAS1* haplotype is one of the clearest examples of adaptive introgression in humans and has undergone strong positive selection in Tibetans (*17, 22, 23*). Beyond *EPAS1*, genome-wide studies have identified multiple loci contributing to Tibetan high-altitude adaptation across diverse physiological systems (*24–32*), and recent work has implicated additional archaic-derived variation in high-altitude–related traits (*33*). Yet it remains unclear whether these signals form a broader and reproducible architecture of adaptive introgression, and whether contributions from different archaic ancestries are organized in similar or distinct ways. The coexistence of Denisovan- and Neanderthal-derived ancestry in Tibetan genomes therefore provides a natural system for addressing these questions.

Here, we analyze whole-genome sequences from 1,039 modern Tibetans (*22, 34*), together with Han Chinese genomes, high-coverage Neanderthal and Denisovan genomes, and temporally stratified ancient genomes from the Tibetan Plateau (*35, 36*). By integrating population-genomic, temporal, environmental and functional evidence, we examine how adaptive introgression from the two archaic ancestries is organized across Tibetan genomes and among individuals. Our results reveal two contrasting architectures—a focal Denisovan contribution dominated by *EPAS1* and a distributed Neanderthal contribution in which heterogeneous introgressed loci converge on shared biological functions—showing that complex adaptation can draw on archaic ancestry without requiring a single shared large-effect haplotype.

### A replicated catalogue reveals contrasting architectures of Tibetan adaptive introgression

We generated independent maps of Tibetan-specific adaptive introgression in TIB1K and TIB38 using the same IBDmix-based framework (*14, 37*) (Fig. 1A, Methods). Despite their different sequencing characteristics, the two datasets yielded highly concordant introgression landscapes and identified 99 Neanderthal-derived and seven Denisovan-derived adaptive haplotypes shared between cohorts (Fig. 1B, figs. S1 to S4 and tables S1 to S2). We therefore used these replicated signals to examine how adaptive introgression from the two archaic ancestries is organized across Tibetan genomes.

**Fig. 1.**
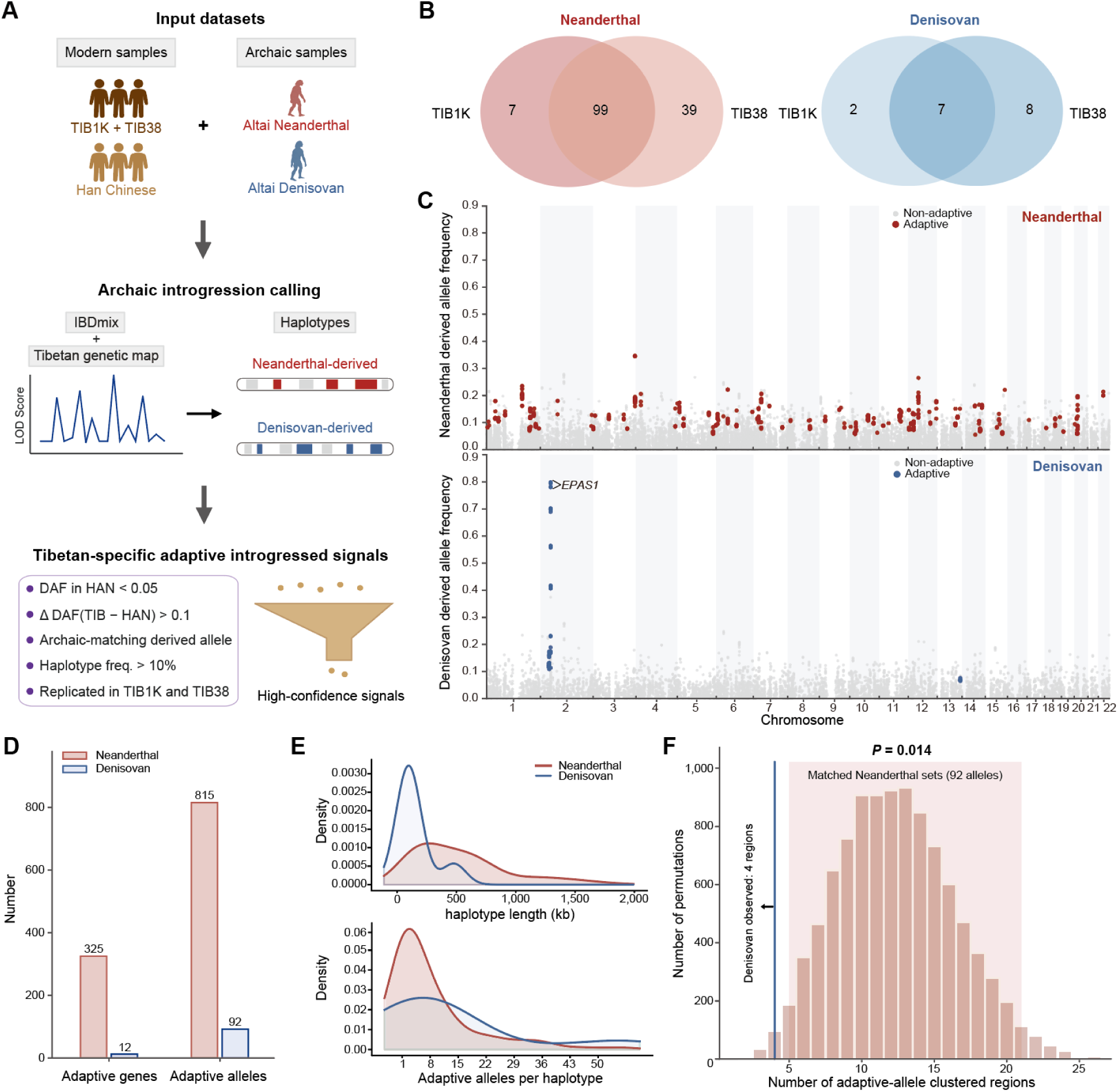
Genome-wide identification of Tibetan-specific adaptive introgressed signals reveals contrasting Neanderthal and Denisovan architectures. **(A)** IBDmix-based identification framework integrating modern and archaic genomes with a Tibetan genetic map. Candidate adaptive introgressed alleles were archaic-matching derived alleles rare in Han Chinese, enriched in Tibetans, within haplotypes carried by >10% of Tibetans, and replicated across TIB1K and TIB38. **(B)** Overlap of independently identified adaptive introgressed haplotypes between cohorts. **(C)** Genome-wide derived allele frequencies for Neanderthal (upper) and Denisovan (lower) ancestry. Colored and grey points indicate adaptive and non-adaptive archaic-matching alleles, respectively. **(D)** Numbers of adaptive introgressed genes and alleles by archaic ancestry. **(E)** Kernel density distributions of haplotype length (upper) and adaptive allele count per haplotype (lower). **(F)** Permutation test assessing whether the stronger genomic clustering of Denisovan-derived adaptive alleles could be explained by their smaller number. Random sets of 92 Neanderthal-derived adaptive alleles, matching the Denisovan count, were grouped into regions using a 200-kb criterion. The histogram shows the resulting null distribution. Denisovan alleles occupied only four regions (blue line), fewer than expected under Neanderthal null distribution (*P* = 0.014). Shading indicates the null distribution’s 2.5th–97.5th percentiles. DAF, derived allele frequency; TIB, Tibetan; HAN, Han Chinese.

The catalogue revealed marked differences in the scale and genomic organization of Neanderthal- and Denisovan-derived adaptive signals. Neanderthal-derived haplotypes contained 815 adaptive alleles across 325 genes, whereas Denisovan-derived haplotypes contained 92 adaptive alleles across 12 genes (Fig. 1, C and D, fig. S4 and tables S3 to S5). Thus, in contrast to the classic emphasis on the Denisovan-derived *EPAS1* locus, the replicated catalogue uncovered a much broader Neanderthal-derived component of Tibetan-specific adaptive introgression (table S3).

The two archaic components also differed in their genome-wide organization. Neanderthal-derived adaptive alleles were distributed across many chromosomes and loci, whereas Denisovan-derived adaptive alleles were concentrated primarily on chromosome 2 near *EPAS1* (Fig. 1C and tables S4 and S5). At the haplotype level, Neanderthal-derived adaptive haplotypes tended to span longer genomic intervals but carried relatively few adaptive alleles per haplotype, consistent with a broad but sparse architecture. Denisovan-derived signals, by contrast, were fewer and more localized, but were locally allele-rich, with multiple adaptive alleles concentrated within a small number of haplotype backgrounds (Fig. 1E). Because Denisovan ancestry is present at much lower genome-wide levels than Neanderthal ancestry in Tibetans, we next wonder whether this apparent clustering could be explained simply by the smaller number of Denisovan-derived adaptive alleles. To test this, we collapsed neighboring adaptive alleles located within 200 kb of one another into single clusters and compared the observed number of Denisovan clusters with matched random sets of Neanderthal-derived adaptive alleles of the same size. Denisovan-derived adaptive alleles resolved into only four clusters, significantly fewer than expected under this matched Neanderthal-derived null distribution, indicating their strong localization is not merely a consequence of the smaller Denisovan-derived allele set but supports stronger clustering of the Denisovan-derived candidate adaptive signals (*P* = 0.014; Fig. 1F).

Together, these results establish a replicated catalogue of Tibetan-specific adaptive archaic introgression and reveal a fundamental architectural contrast: a larger and more dispersed Neanderthal-derived component, and a smaller but highly localized Denisovan-derived component dominated by the *EPAS1* region.

### Temporal and elevational dynamics reinforce contrasting archaic architectures

We next asked whether the contrasting genomic architectures identified in present-day Tibetans were also reflected in allele-frequency changes across temporal and environmental gradients. To examine temporal dynamics, we analyzed 156 ancient individuals from the Tibetan Plateau spanning approximately 5,100 to 700 years before present (*35, 36*). These individuals represented six broad geographic regions of the Plateau and were grouped into four chronological intervals according to their radiocarbon dates (Fig. 2A). After filtering for sufficient callability in the ancient genomes, 79 Neanderthal-derived and 13 Denisovan-derived adaptive alleles were retained for analysis (tables S6 and S7).

**Fig. 2.**
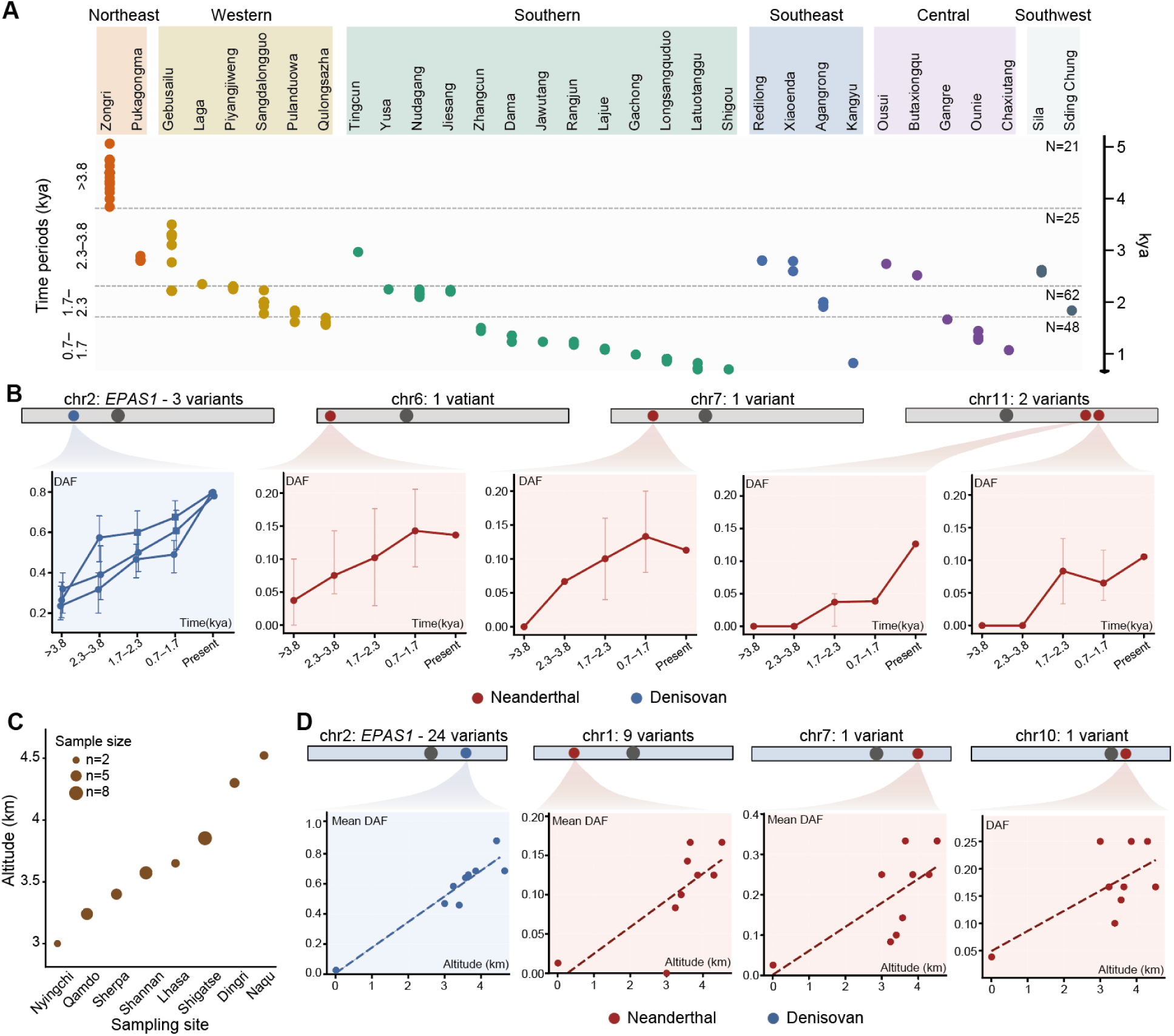
Temporal and elevational dynamics reinforce contrasting archaic architectures. **(A)** Geographic and temporal distribution of 156 ancient Tibetan Plateau individuals. Individuals are arranged by archaeological site within six broad geographic regions, indicated by colors. Each point represents one individual. Dashed lines separate four chronological intervals spanning approximately 5.1–0.7 thousand years ago (kya); *N* indicates sample size per interva*l*. (**B)** Temporal trajectories of adaptive archaic alleles showing significant increases in frequency towards the present. Points indicate derived allele frequencies across four ancient intervals and present-day Tibetans; error bars indicate 95% confidence intervals. Chromosome schematics mark allele positions. Red and blue denote Neanderthal- and Denisovan-derived alleles, respectively. (**C)** Elevations and sample sizes of eight Tibetan groups from TIB38. Sites are ordered by elevation; point heights indicate elevation and sizes represent sample counts. (**D)** Elevational trajectories of adaptive archaic alleles showing significant positive correlations between derived allele frequency and altitude. Chromosome schematics mark introgressed regions. Points show group-specific mean derived allele frequencies across 24 Denisovan-derived alleles at *EPAS1* region (chr2) and nine Neanderthal-derived alleles on chr1, or individual variant frequencies for chr7 and chr10. Dashed lines indicate linear regression fits. Red and blue denote Neanderthal- and Denisovan-derived alleles, respectively.

Among these alleles, four Neanderthal-derived variants showed significant increases in frequency towards the present (Pearson’s *r* > 0.8, *P* < 0.05). These variants occurred at multiple genomic locations, including regions on chromosomes 6, 7 and 11 (Fig. 2B and table S6). By contrast, all three Denisovan-derived variants showing significant temporal increases were intronic variants within *EPAS1* on chromosome 2 (Pearson’s *r* > 0.9, *P* < 0.05; Fig. 2B and table S7). Thus, although temporally dynamic variants were detected for both archaic ancestries, the Denisovan signal was confined to the canonical *EPAS1* locus, whereas the Neanderthal signals occurred across several independent genomic regions.

We next examined adaptive archaic allele frequencies varied across present-day Tibetan populations sampled at an elevational gradient. The TIB38 dataset^34^ comprised individuals from eight locations ranging from approximately 3,000 to 4,500 m above sea level (Fig. 2C). Eleven Neanderthal-derived adaptive alleles showed positive correlations between allele frequency and altitude (Pearson’s *r* > 0.6, *P* < 0.05), forming three genomic clusters on chromosomes 1, 7 and 10 (Fig. 2D and table S8). In contrast, 24 Denisovan-derived alleles were significantly correlated with altitude, all of which fell within the *EPAS1*-centered introgressed region on chromosome 2 (Pearson’s *r* > 0.7, *P* < 0.05; Fig. 2D and table S9). Notably, the three *EPAS1* variants that increased in frequency through time were also among those positively correlated with altitude, providing concordant temporal and elevational evidence at the same Denisovan-derived locus.

Together, these two complementary analyses reinforce the architectural contrast observed in the genome-wide catalogue. Across both temporal and elevational dimensions, dynamic Denisovan-derived signals repeatedly converged on the *EPAS1* region, whereas Neanderthal-derived candidate signals occurred across multiple genomic regions. These patterns highlight an *EPAS1*-dominated Denisovan trajectory while supporting a more spatially distributed contribution from Neanderthal-derived adaptive variation.

### Adaptive archaic alleles are predominantly noncoding and occupy regulatory genomic contexts

Having established contrasting genomic and evolutionary patterns of Tibetan-specific adaptive archaic introgression, we next examined the potential functional contexts of these variants. Gene Ontology annotations (*38*) of genes carrying adaptive introgressed alleles implicated a broad range of biological processes, including developmental processes, vesicle-transport and neurodevelopmental functions for Neanderthal-derived genes and hypoxia-related, calcium-homeostasis and transcriptional processes for Denisovan-derived genes (fig. S5, tables S10 and S11). We therefore focused on the molecular consequences and regulatory contexts of the underlying adaptive alleles.

Variant Effect Predictor (*39*) annotation showed that the candidate adaptive alleles were overwhelmingly noncoding. Of 815 Neanderthal-derived adaptive alleles, 813 occurred outside protein-coding sequences, and all 92 Denisovan-derived adaptive alleles were noncoding (Fig. 3A and tables S4 and S5). Only two Neanderthal-derived alleles altered coding sequence, including the missense variant rs2289491-C in *SLC9A9*, a sodium/proton exchanger gene linked to neurological traits (*40*). This strong predominance of noncoding variation prompted us to test whether adaptive archaic alleles were preferentially located in regulatory genomic elements.

**Fig. 3.**
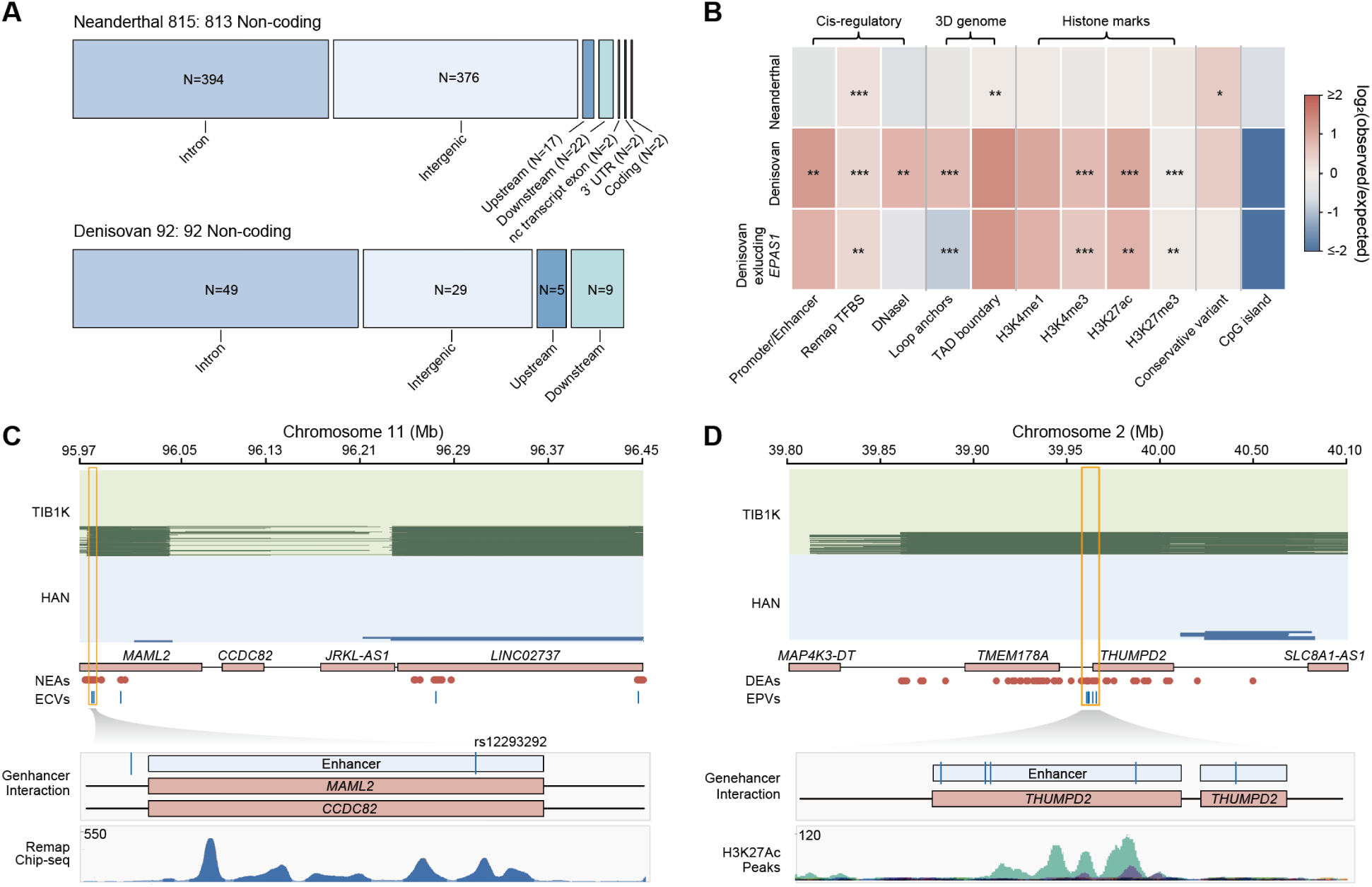
Adaptive archaic alleles are predominantly noncoding and show regulatory enrichment. **(A)** Variant consequences of Neanderthal- and Denisovan-derived adaptive alleles annotated using the Variant Effect Predictor. (**B)** Permutation-based enrichment of adaptive archaic alleles across regulatory, chromatin and evolutionary-conservation annotations. Rows represent Neanderthal-derived alleles, all Denisovan-derived alleles and Denisovan-derived alleles after excluding the *EPAS1*-centred introgressed regions. Color indicates log_2_(observed/expected), where expected values are derived from permutations. Red and blue denote enrichment and depletion, respectively; color values are capped at −2 and 2. Asterisks indicate significance from empirical permutation tests. (**C)** A Neanderthal-derived introgressed region on chr11 near *MAML2*, showing introgressed segments in TIB1K and Han Chinese individuals, Neanderthal-derived adaptive alleles (NEAs) and evolutionarily conserved variants (ECVs). The highlighted interval is expanded below to show GeneHancer enhancer–gene interactions and ReMap ChIP–seq peaks. (**D)** A Denisovan-derived introgressed region on chr2 near *THUMPD2*, showing introgressed segments in TIB1K and Han Chinese individuals, Denisovan-derived adaptive alleles (DEAs) and enhancer/promoter variants (EPVs). The highlighted interval is expanded below to show GeneHancer enhancer–gene interactions and H3K27ac peaks.

We therefore performed permutation-based enrichment analyses across promoter and enhancer elements (*41*), evolutionary conservation metrics (*42, 43*), transcription factor binding sites (*44*), histone modifications (*45*), CpG islands (*46*), chromatin interaction features (*47*), and DNase I hypersensitivity sites (*45*). Adaptive alleles from both archaic ancestries were significantly enriched in transcription factor binding sites (*P* < 0.001; Fig. 3B). Neanderthal-derived alleles were additionally enriched in evolutionarily conserved regions (*P* = 0.0012) and regions marked by H3K27me3 (*P* = 0.018). Denisovan-derived alleles were enriched in promoter and enhancer elements (*P* = 0.007), TAD boundaries (*P* < 0.001), loop anchors (*P* = 0.005) and the active histone marks H3K4me1, H3K4me3 and H3K27ac (all *P* < 0.001). Because a large proportion of Denisovan-derived adaptive alleles cluster around *EPAS1*, we repeated the analysis after excluding the *EPAS1*-centered introgressed region. The remaining Denisovan-derived alleles retained significant enrichment in transcription factor binding sites, TAD boundaries and the active histone marks H3K4me1, H3K4me3 and H3K27ac (Fig. 3B), indicating that these regulatory enrichments were not driven solely by the canonical *EPAS1* locus.

Representative loci illustrate these regulatory contexts at individual introgressed regions (Fig. 3, C and D). At the *MAML2* locus on chromosome 11, a Tibetan-specific Neanderthal-derived segment contains 28 consecutive adaptive alleles, five of which fall within highly conserved sequence. One of these variants, rs12293292-T, overlaps an enhancer linked to *MAML2*, a regulator of cell proliferation and differentiation (*48*), and *CCDC82*, a gene associated with neurodevelopmental traits (*49*), and also overlaps transcription factor binding peaks from ReMap, together supporting plausible cis-regulatory activity at this locus (Fig. 3C). At the *THUMPD2* locus on chromosome 2, outside the canonical *EPAS1* region, a Denisovan-derived segment contains 55 consecutive adaptive alleles, five of which overlap an H3K27ac-marked enhancer linked to *THUMPD2*, a gene implicated in retinal function (*50*) (Fig. 3D). These examples further illustrate how Tibetan-specific adaptive archaic variation is embedded within regulatory genomic contexts beyond the canonical *EPAS1* locus.

We next evaluated whether subsets of these noncoding adaptive alleles were linked to gene expression or phenotypic variation in existing functional datasets. GTEx annotations (*51*) identified 153 Neanderthal-derived adaptive alleles as expression quantitative trait loci (eQTLs), corresponding to 72 eGenes across 23 broad tissue categories (fig. S6A), including esophagus, skin, adipose tissue, thyroid, nerve, heart, brain, muscle and blood (fig. S6B). Fourteen Denisovan-derived adaptive alleles were annotated as eQTLs, corresponding to seven eGenes across six tissue categories. Integration with the Tibetan-specific HiLand GWAS resource (*52*) further identified 90 Neanderthal-derived adaptive alleles associated at a nominal threshold of *P* < 0.001 with 39 traits across six broad categories (fig. S6C), including hematological, metabolic, ocular, anthropometric, cardiac and reproductive traits (fig. S6D and table S12). Nineteen Denisovan-derived adaptive alleles were associated with eight traits across four categories, concentrated in hematological, metabolic, anthropometric and reproductive traits (fig. S6D and table S13). Five Denisovan-derived alleles reached genome-wide significance (*P* < 5 × 10⁻⁸), for altitude- and red-blood-cell-related traits (table S13).

Together, these analyses show that Tibetan-specific adaptive archaic alleles are overwhelmingly noncoding and are preferentially located in regulatory genomic contexts. Their enrichment patterns span transcription factor binding, chromatin state and three-dimensional genome organization, with regulatory signatures persisting for Denisovan-derived alleles even outside *EPAS1*. Existing eQTL and Tibetan GWAS resources further link subsets of these variants to gene-expression and phenotypic variation, providing functional context for the broader adaptive architectures identified above.

### Heterogeneous Neanderthal introgressed repertoires show recurrent functional convergence

Having characterized population-level adaptive introgression, we next asked how archaic-derived variation is organized within individual Tibetan genomes. The high-confidence adaptive signals identified above represent only a subset of the introgressed genes carried by each individual (fig. S7). We therefore examined the full repertoire of archaic introgressed genes in each TIB1K genome. Pairwise comparisons revealed substantial individual-level heterogeneity, with mean overlap proportions of only 11.55% for Neanderthal-derived and 14.96% for Denisovan-derived gene repertoires (Fig. 4A). We therefore asked whether these heterogeneous introgressed repertoires nevertheless recurrently mapped to shared biological functions.

**Fig. 4.**
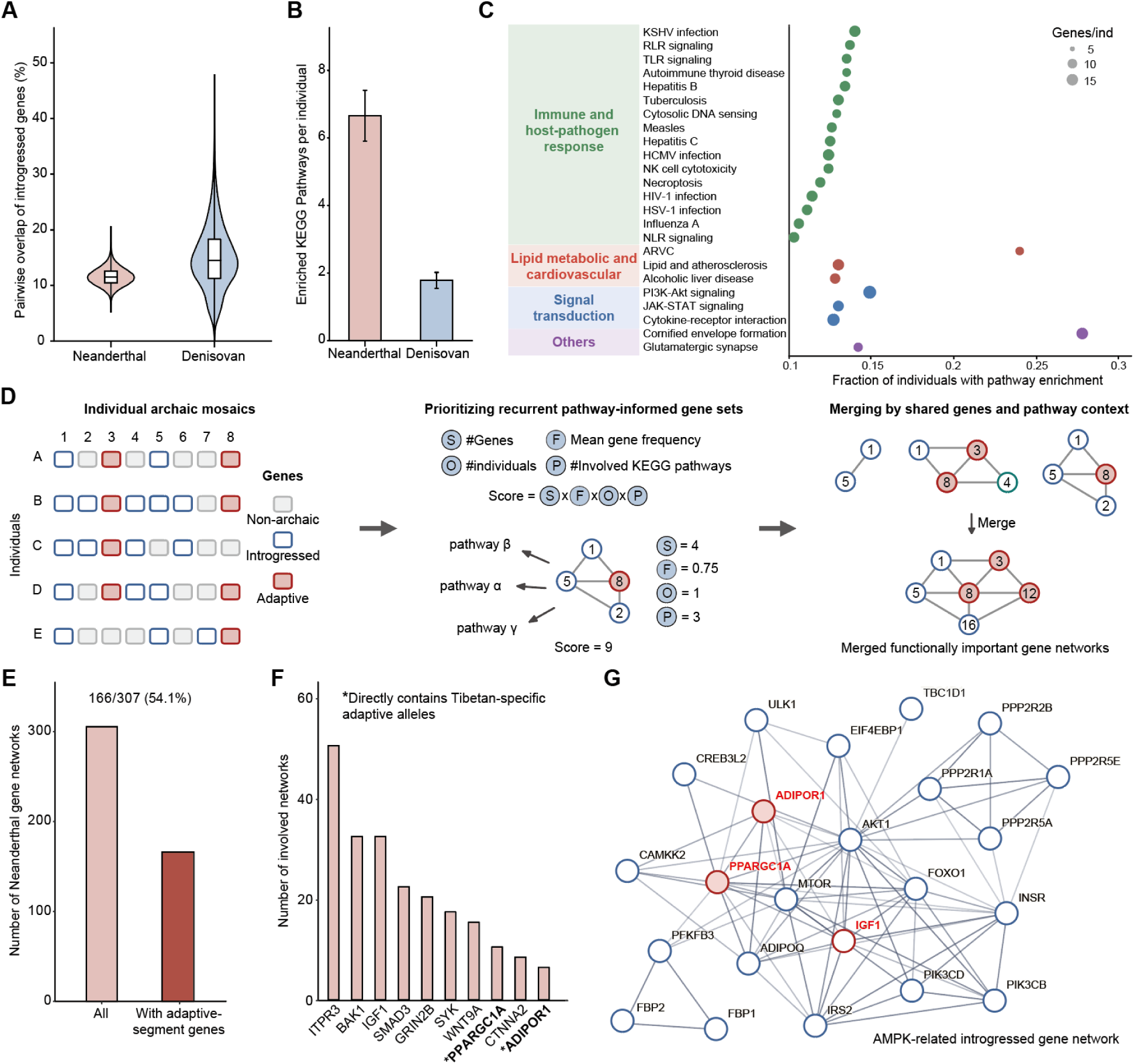
Heterogeneous Neanderthal introgressed repertoires show recurrent functional convergence. **(A)** Pairwise overlap of introgressed genes between TIB1K individuals by archaic ancestry. Boxes show medians and interquartile ranges. (**B)** Numbers of enriched KEGG pathways per individual, shown as means with 95% confidence intervals. (**C)** Recurrently enriched Neanderthal-derived KEGG pathways grouped by functional category. Horizontal positions indicate enrichment prevalence across individuals; point sizes indicate introgressed gene counts per individual, and colors denote functional categories. (**D)** Schematic of the framework used to identify functionally important introgressed gene networks recurrently shared across individuals. For each individual, KEGG-defined gene sets were scored by introgressed gene count, mean gene frequency, carrier count, and pathway count. Top-ranked sets were merged through shared genes within pathway contexts. (**E)** Number of Neanderthal-derived gene networks containing genes that overlap Tibetan-specific adaptive introgressed segments (166/307; 54.1%). (**F)** Adaptive-segment genes recurring across Neanderthal-derived networks. Bars indicate network counts; asterisks identify genes directly containing Tibetan-specific adaptive introgressed alleles. (**G)** Representative AMPK-related Neanderthal-derived pathway-informed network. Red outlines and labels indicate genes overlapping Tibetan-specific adaptive introgressed segments; light red fill identifies genes directly containing adaptive alleles. White nodes with blue outlines indicate other Neanderthal-introgressed genes.

For each individual, we performed KEGG (*53*) pathway enrichment using the full set of introgressed genes carried in that genome. Neanderthal-derived gene repertoires yielded an average of 6.66 enriched pathways per individual, compared with 1.79 for Denisovan-derived repertoires (Fig. 4B). We next defined recurrent pathways as those enriched in more than 10% of TIB1K individuals. Twenty-four Neanderthal-derived pathways met this criterion, compared with one Denisovan-derived pathway. Because the Neanderthal-derived introgressed gene repertoire is substantially larger than the Denisovan-derived repertoire, we did not interpret these absolute pathway counts as evidence for stronger functional convergence of Neanderthal ancestry. Instead, we focused on whether heterogeneous Neanderthal-derived repertoires themselves showed recurrent functional organization across individuals. The recurrent Neanderthal-derived pathways fell into five broad functional modules encompassing immune and host–pathogen response, lipid metabolic and cardiovascular biology, signal transduction, epithelial barrier function and neuronal signaling (Fig. 4C). Thus, despite limited sharing of individual introgressed genes, Neanderthal-derived repertoires repeatedly mapped to a common set of biological processes across Tibetan genomes.

We next asked whether this pathway-level recurrence could be resolved into recurrent combinations of introgressed genes. We developed a pathway-informed scoring framework that prioritizes gene sets on the basis of their size, mean introgression frequency, recurrence across individuals and involvement across KEGG pathways (Fig. 4D). Gene sets containing at least two introgressed genes within a common pathway were scored using these four features; the top 5% were retained and subsequently merged according to shared genes and pathway context to generate prioritized pathway-informed introgressed gene networks. This analysis identified 307 Neanderthal-derived gene networks, 166 of which contained at least one Tibetan-specific adaptive introgressed gene (Fig. 4E and table S14).

Adaptive-segment genes were unevenly represented across these networks. Several genes located within Tibetan-specific adaptive introgressed segments occurred repeatedly in different network contexts, linking population-level adaptive candidates to recurrent individual-level functional modules (Fig. 4F and table S14). Among these genes, *PPARGC1A* and *ADIPOR1* also directly contained Tibetan-specific adaptive introgressed alleles. A representative AMPK-related pathway-informed network included *PPARGC1A*, *ADIPOR1*, *IGF1* and other genes within Neanderthal-derived adaptive introgressed regions, together with additional Neanderthal-introgressed genes, linking this recurrent network to energy sensing, lipid metabolism, glucose homeostasis and insulin-related biology (Fig. 4G and table S14).

Together, these analyses reveal a notable feature of the Neanderthal-derived component of Tibetan genomes: although individuals carry markedly different combinations of introgressed genes, these heterogeneous repertoires repeatedly map to a restricted set of biological pathways and pathway-informed gene networks. This recurrent functional organization provides an individual-genome perspective on the distributed Neanderthal-derived architecture identified at the population level and nominates metabolically relevant adaptive genes, including *PPARGC1A*, for further phenotypic investigation.

### A Tibetan-enriched Neanderthal *PPARGC1A* haplotype links adaptive introgression to glycemic variation

Among genes directly carrying Tibetan-specific adaptive introgressed alleles, *PPARGC1A* emerged as a particularly prominent candidate, being the most recurrently represented across the prioritized Neanderthal-derived pathway-informed networks. *PPARGC1A* encodes a transcriptional coactivator with central roles in energy metabolism, mitochondrial biogenesis, lipid metabolism and glucose homeostasis, and has been extensively implicated in type 2 diabetes (T2D)-related biology (*54, 55*). At this locus, we identified three Tibetan-specific adaptive introgressed alleles within a Neanderthal-derived introgressed segment. This introgressed segment was substantially more frequently carried in TIB1K than in the Han Chinese comparison panel, being present in 311 of 1,001 TIB1K individuals and 3 of 39 Han Chinese individuals (Fig. 5A). We then extended the three adaptive alleles using high-linkage-disequilibrium sites (*r*² > 0.8) carrying the Neanderthal-matching allele, defining a seven-variant Neanderthal-derived haplotype (Methods). Comparison of TIB1K with 1KGP populations revealed marked population differentiation in the frequency of this haplotype: its allele frequency reached approximately 17% in TIB1K, compared with 1–12% in other Asian populations, and it was undetected in the sampled populations outside Asia (Fig. 5B). Thus, both segment-level carriage and haplotype-frequency comparisons indicate enrichment of Neanderthal-derived variation at the *PPARGC1A* locus in Tibetans.

**Fig. 5.**
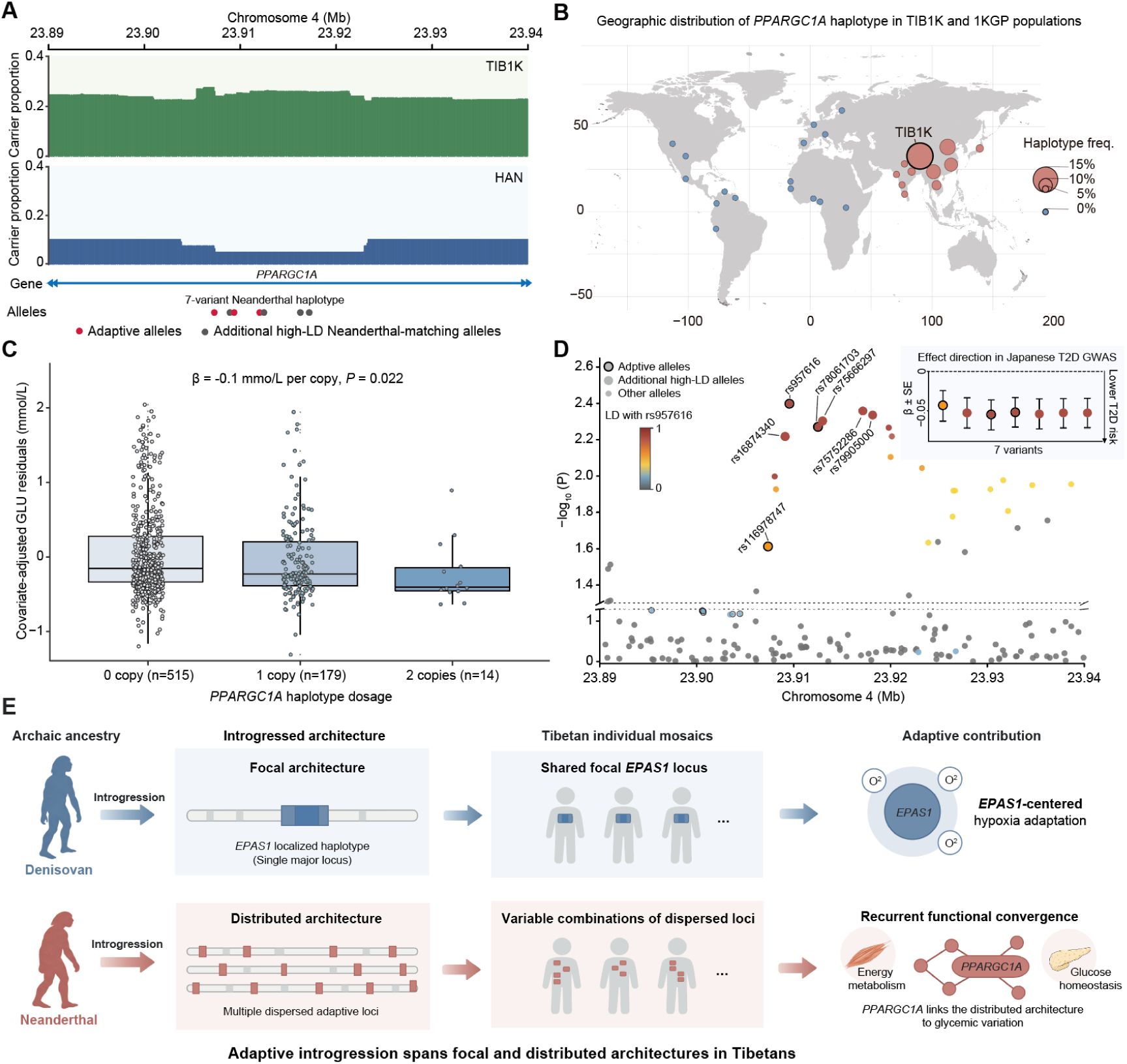
A Tibetan-enriched Neanderthal *PPARGC1A* haplotype links adaptive introgression to glycemic variation. **(A)** Introgressed-segment carrier proportions across *PPARGC1A* in TIB1K (green; n = 1,001) and Han Chinese (blue; n = 39). Three adaptive alleles (red) and four additional Neanderthal-matching variants (grey; r² > 0.8) define the haplotype. (**B)** Haplotype distribution in TIB1K and 1KGP populations. Circle sizes indicate haplotype allele frequencies; red and blue denote nonzero and zero frequencies, respectively. (**C)** Covariate-adjusted fasting glucose residuals by haplotype dosage (0/1/2 copies; n = 515/179/14). The additive model estimates the glucose change per haplotype copy (β, mmol/L) and its *P* value. (**D)** Japanese T2D GWAS associations from GWAS Atlas, colored by LD with the lead SNP rs957616. Enlarged points indicate the seven haplotype variants; black outlines identify adaptive alleles. Inset: effects (*β* ± SE) aligned to Neanderthal-derived alleles and ordered by genomic position. Negative effects indicate lower T2D risk. (**E)** Conceptual model contrasting focal Denisovan contributions centered on *EPAS1* and hypoxia adaptation with dispersed Neanderthal loci showing recurrent functional convergence across individuals. *PPARGC1A* exemplifies the latter architecture through associations with glycemic variation.

This population pattern was particularly notable in light of the metabolic epidemiology of Tibetan populations (*56*). Large population-based studies have reported a substantially lower prevalence of diabetes in Tibetans than in Han Chinese (*57, 58*), while diabetes risk within Tibetan populations has also been reported to decrease with increasing altitude (*59*). These observations raise the possibility that genetic changes associated with long-term high-altitude adaptation may intersect with pathways influencing metabolic physiology and diabetes susceptibility. Given the central role of *PPARGC1A* in glucose and energy metabolism, we therefore asked whether the introgressed haplotype was associated with fasting glucose variation in Tibetans.

Fasting glucose measurements and complete covariate information were available for 708 TIB1K individuals included in the association analysis. Because the seven Neanderthal-derived variants are in near-complete linkage disequilibrium and represent the same haplotype background, we tested the dosage of the introgressed haplotype rather than individual variants. Each additional copy of the *PPARGC1A* introgressed haplotype was associated with a 0.10 mmol/L reduction in fasting glucose (*β* = −0.1, *P* = 0.022; Fig. 5C and table S15). Although modest in magnitude, this association provides phenotypic evidence linking a Tibetan-enriched adaptive introgressed haplotype to fasting-glucose variation.

We then asked whether variation at the same locus showed independent evidence of association with T2D. In a Japanese population-based T2D GWAS (*60*), the seven variants defining the introgressed haplotype coincided with the strongest association signals across the *PPARGC1A* introgressed region, and the Neanderthal-derived alleles consistently showed effect directions associated with reduced T2D risk (Fig. 5D and table S16). This direction was concordant with the lower fasting glucose associated with increasing haplotype dosage in TIB1K. Although the Japanese GWAS does not directly replicate the Tibetan haplotype association, it provides independent support linking variation at the same introgressed locus to T2D risk in another East Asian population.

Together, these results provide convergent evidence linking the Neanderthal-derived *PPARGC1A* haplotype to metabolic variation. The haplotype is enriched in Tibetans, associated with lower fasting glucose in TIB1K, and supported by directionally concordant T2D association signals in an independent East Asian population. These findings nominate *PPARGC1A* as a trait-relevant candidate within the distributed Neanderthal-derived adaptive architecture.

The *PPARGC1A* locus also provides a trait-relevant example of the broader architecture identified in our study (Fig. 5E). Whereas the Denisovan contribution to Tibetan adaptation is dominated by a focal *EPAS1*-centered architecture associated with hypoxia adaptation, Neanderthal-derived adaptive variation is distributed across multiple loci and occurs in different combinations among individuals, yet repeatedly converges on shared biological functions. *PPARGC1A* illustrates how one component of this distributed architecture can be connected to measurable glycemic variation, supporting a broader model in which adaptive introgression in Tibetans spans focal and distributed genomic architectures.

## Discussion

Adaptive introgression is often conceptualized through individual loci of large effect, with the Denisovan-derived *EPAS1* haplotype in Tibetans representing one of its clearest examples (*17,22, 23*). Our results reveal that this focal architecture represents only one way in which archaic ancestry can contribute to human adaptation. In Tibetan genomes, Denisovan-derived adaptive variation is dominated by a highly localized *EPAS1*-centred component that is widely shared across individuals, whereas the Neanderthal-derived component is distributed across multiple genomic regions and occurs in different combinations among individuals. Yet these heterogeneous Neanderthal-derived repertoires repeatedly converge on related biological functions. We therefore propose that adaptive introgression in Tibetans spans focal and distributed architectures: one organized around a shared, strongly selected archaic haplotype, and another in which multiple introgressed loci are distributed across the genome but show recurrent functional convergence. These architectures should not be viewed as intrinsic properties of Denisovan versus Neanderthal ancestry; rather, they likely reflect the combined effects of ancestry abundance, introgression history, recombination, selection strength and the genetic architecture of the traits under selection.

Several features of the data support this architectural contrast. The distributed Neanderthal component was reproducibly detected in two independently analyzed Tibetan datasets, arguing against its being driven by a single cohort or sequencing design. Importantly, the strong localization of Denisovan-derived adaptive variation was not simply a consequence of the smaller number of Denisovan-derived candidate alleles: even after matching the number of Neanderthal-derived alleles, Denisovan signals occupied significantly fewer genomic regions. The same contrast was also evident across two complementary evolutionary dimensions. Adaptive alleles showing temporal frequency changes in ancient Tibetan Plateau genomes and those correlated with elevation in contemporary Tibetans repeatedly converged on *EPAS1* for Denisovan ancestry, whereas the corresponding Neanderthal-derived signals occurred across multiple loci. These temporal and elevational analyses therefore provide orthogonal support for the genomic architecture identified in present-day populations, although they should not be interpreted as independent demonstrations of selection at every candidate locus. Allele-frequency trajectories can also be influenced by drift, demographic structure, local population history and heterogeneous sampling, particularly given the still-limited temporal and geographic resolution of ancient Tibetan genomes.

A shared feature of both architectures is the overwhelming predominance of noncoding adaptive variation, pointing to gene regulation as an important route through which archaic ancestry may influence Tibetan adaptation. Both Neanderthal-and Denisovan-derived adaptive alleles were enriched in regulatory genomic elements, although their enrichment profiles pointed to partly different regulatory contexts. Neanderthal-derived alleles were associated with conserved sequence and repressive chromatin annotations, whereas Denisovan-derived alleles were enriched in promoter–enhancer, active chromatin and three-dimensional genome annotations. Notably, several Denisovan regulatory enrichments persisted after removal of the *EPAS1* region, indicating that their regulatory signature is not attributable solely to this canonical locus. These patterns should not be interpreted as demonstrating fundamentally different molecular mechanisms between the two ancestries, because direct statistical comparisons of enrichment magnitude were not performed. Rather, they indicate that adaptive archaic variation from both sources has been preferentially retained in regulatory contexts, with potentially different distributions across regulatory features. More complete Tibetan-specific transcriptomic, epigenomic and single-cell datasets will be important for resolving the tissue specificity, direction and biological consequences of these variants.

The individual-genome analyses provide a complementary perspective on what a distributed adaptive architecture may entail. Individual Tibetans share only a limited fraction of their exact introgressed gene repertoires, reflecting the distinct mosaics created by recombination, inheritance and demographic history. Despite this heterogeneity, Neanderthal-derived repertoires repeatedly mapped to a restricted set of immune, metabolic and signal-transduction pathways, and adaptive introgressed genes recurred within pathway-informed multi-gene networks. Thus, functional similarity need not require individuals to share the same introgressed loci: different combinations of archaic genes can repeatedly map onto related biological programs. This organization differs conceptually from the focal *EPAS1* model, in which the same strongly selected haplotype is shared by a large fraction of the population. At the same time, we do not equate this distributed architecture with classical polygenic adaptation. Our analyses do not demonstrate coordinated selection on a defined polygenic trait or quantify the joint phenotypic effects of multiple introgressed loci. Rather, they show that multiple Neanderthal-derived adaptive candidates, variably represented among individuals, converge at the level of biological function. Because the pathway and network analyses are informed by existing KEGG annotations and the Neanderthal-derived gene repertoire is substantially larger than the Denisovan-derived repertoire, they are best viewed as a framework for identifying recurrent functional organization and generating experimentally testable hypotheses rather than as evidence for de novo interaction networks.

The *PPARGC1A* locus provides a trait-relevant example of how such a distributed architecture may intersect with contemporary complex phenotypes. *PPARGC1A* emerged from the recurrent Neanderthal-derived metabolic networks and directly contains Tibetan-specific adaptive introgressed alleles. Neanderthal-derived variation at this locus is enriched in Tibetans, and increasing dosage of the seven-variant introgressed haplotype was nominally associated with lower fasting glucose, with an estimated reduction of 0.10 mmol/L per haplotype copy. Variation within the same introgressed region also showed directionally concordant associations with reduced T2D risk in an independent Japanese GWAS (*60*). This convergence is particularly intriguing in light of epidemiological studies reporting lower diabetes prevalence in Tibetans than in Han Chinese and lower diabetes risk with increasing altitude among Tibetans (*57–59*). We do not infer that the *PPARGC1A* haplotype explains these population-level epidemiological differences, nor does the Japanese GWAS constitute replication of the Tibetan haplotype association. The Tibetan glucose association is modest and nominal, and the available phenotype analysis was restricted to pregnant Tibetan women (*34*), which further limits its generalizability. Nevertheless, the agreement among evolutionary enrichment, a glycemic phenotype in Tibetans and independent T2D genetic evidence raises the possibility that archaic variants retained during high-altitude adaptation can have pleiotropic consequences for present-day metabolic physiology. Replication in broader Tibetan cohorts across sexes and physiological states, together with functional characterization of the haplotype, will be required to establish its metabolic effects and causal relevance.

More broadly, our study expands the scale at which adaptive introgression can be considered. The resolution of the temporal analyses remains constrained by ancient-DNA availability, most functional genomic resources are derived from non-Tibetan populations, and future analyses using more complete telomere-to-telomere references may further refine the underlying introgressed landscape (*61, 62*). These limitations notwithstanding, the combined genomic, temporal, regulatory and individual-level evidence reveals an organizational contrast that is not apparent from the canonical *EPAS1* example alone. A focal archaic haplotype can provide a powerful route to adaptation, but adaptive introgression can also be distributed across loci that differ among individuals while converging on related biological functions. Adaptive introgression in Tibetans therefore spans focal and distributed architectures, linking archaic evolutionary history not only to adaptation in an extreme environment but also to present-day complex-trait variation.

## Acknowledgments

We thank all members of ChenLab at Fudan University for their support and insightful discussions related to this work. We also thank J. Spence for his assistance with generating the genetic map of Tibetans. Computational resources were provided by the CFFF High-Performance Computing Platform at Fudan University.

## Author contributions

Conceptualization: LC, SX, YH Methodology: SL, QZ, XG, YX, KL, ZX, QF Investigation: SL, QZ, XG

Visualization: SL, QZ Funding acquisition: LC Project administration: LC Supervision: LC, SX, YH

Writing – original draft: SL, QZ, LC

Writing – review & editing: XG, YX, KL, ZX, LD, XG, BS, QF, YZ, YH, SX

## Data, code, and materials availability

Previously published whole-genome sequencing data for the TIB1K cohort are available from the Genome Sequence Archive for Human (GSA-Human) under BioProject accession PRJCA007843 and study accession HRA001809 (https://bigd.big.ac.cn/gsa-human/browse/HRA001809) (*34*). Whole-genome sequencing data for the TIB38 and Han Chinese cohorts are available from the Genome Sequence Archive under accession PRJCA000246 and the National Omics Data Encyclopedia under accession ND00000013EP (*22*). Sequence data for the Altai Neanderthal and Altai Denisovan genomes are available from the European Nucleotide Archive under accessions ERP002097(*2*) and ERP001519 (*6*), respectively. Tibetan introgression calls generated in this study and the code for network convergence analyses are available from Zenodo at https://doi.org/10.5281/zenodo.22763707 (*63*).

## Supplementary Materials

Materials and Methods

Figs. S1 to S7

Tables S1 to S16

References (*64–67*)

## References and Notes

1. R. E. Green et al., A draft sequence of the Neandertal genome. Science 328, 710–722 (2010).

2. K. Prüfer et al., The complete genome sequence of a Neanderthal from the Altai Mountains. Nature 505, 43–49 (2014).

3. K. Prüfer et al., A high-coverage Neandertal genome from Vindija Cave in Croatia. Science 358, 655–658 (2017).

4. F. Mafessoni et al., A high-coverage Neandertal genome from Chagyrskaya Cave. Proc Natl Acad Sci U S A 117, 15132–15136 (2020).

5. D. Reich et al., Genetic history of an archaic hominin group from Denisova Cave in Siberia. Nature 468, 1053–1060 (2010).

6. M. Meyer et al., A high-coverage genome sequence from an archaic Denisovan individual. Science 338, 222–226 (2012).

7. S. Sankararaman, S. Mallick, N. Patterson, D. Reich, The Combined Landscape of Denisovan and Neanderthal Ancestry in Present-Day Humans. Curr Biol 26, 1241–1247 (2016).

8. B. Vernot et al., Excavating Neandertal and Denisovan DNA from the genomes of Melanesian individuals. Science 352, 235–239 (2016).

9. S. R. Browning, B. L. Browning, Y. Zhou, S. Tucci, J. M. Akey, Analysis of Human Sequence Data Reveals Two Pulses of Archaic Denisovan Admixture. Cell 173, 53–61.e59 (2018).

10. S. Mallick et al., The Simons Genome Diversity Project: 300 genomes from 142 diverse populations. Nature 538, 201–206 (2016).

11. G. S. Jacobs et al., Multiple Deeply Divergent Denisovan Ancestries in Papuans. Cell 177, 1010–1021.e1032 (2019).

12. M. Larena et al., Philippine Ayta possess the highest level of Denisovan ancestry in the world. Curr Biol 31, 4219–4230.e4210 (2021).

13. J. D. Wall et al., Higher Levels of Neanderthal Ancestry in East Asians than in Europeans. Genetics 194, 199-+ (2013).

14. L. Chen, A. B. Wolf, W. Fu, L. Li, J. M. Akey, Identifying and Interpreting Apparent Neanderthal Ancestry in African Individuals. Cell 180, 677–687.e616 (2020).

15. S. Sankararaman et al., The genomic landscape of Neanderthal ancestry in present-day humans. Nature 507, 354–357 (2014).

16. B. Vernot, J. M. Akey, Resurrecting surviving Neandertal lineages from modern human genomes. Science 343, 1017–1021 (2014).

17. E. Huerta-Sánchez et al., Altitude adaptation in Tibetans caused by introgression of Denisovan-like DNA. Nature 512, 194–197 (2014).

18. M. Dannemann, J. Kelso, The Contribution of Neanderthals to Phenotypic Variation in Modern Humans. Am J Hum Genet 101, 578–589 (2017).

19. R. M. Gittelman et al., Archaic Hominin Admixture Facilitated Adaptation to Out-of-Africa Environments. Curr Biol 26, 3375–3382 (2016).

20. F. Racimo et al., Archaic Adaptive Introgression in TBX15/WARS2. Mol Biol Evol 34, 509–524 (2017).

21. C. N. Simonti et al., The phenotypic legacy of admixture between modern humans and Neandertals. Science 351, 737–741 (2016).

22. D. Lu et al., Ancestral Origins and Genetic History of Tibetan Highlanders. Am J Hum Genet 99, 580–594 (2016).

23. H. Hu et al., Evolutionary history of Tibetans inferred from whole-genome sequencing. PLoS Genet 13, e1006675 (2017).

24. X. Yi et al., Sequencing of 50 human exomes reveals adaptation to high altitude. Science 329, 75–78 (2010).

25. Y. Peng et al., Genetic variations in Tibetan populations and high-altitude adaptation at the Himalayas. Mol Biol Evol 28, 1075–1081 (2011).

26. S. Xu et al., A genome-wide search for signals of high-altitude adaptation in Tibetans. Mol Biol Evol 28, 1003–1011 (2011).

27. C. M. Beall et al., Natural selection on EPAS1 (HIF2alpha) associated with low hemoglobin concentration in Tibetan highlanders. Proc Natl Acad Sci U S A 107, 11459–11464 (2010).

28. T. S. Simonson et al., Genetic evidence for high-altitude adaptation in Tibet. Science 329, 72–75 (2010).

29. L. Deng et al., Prioritizing natural-selection signals from the deep-sequencing genomic data suggests multi-variant adaptation in Tibetan highlanders. Natl Sci Rev 6, 1201–1222 (2019).

30. J. Yang et al., Genetic signatures of high-altitude adaptation in Tibetans. Proc Natl Acad Sci U S A 114, 4189–4194 (2017).

31. J. Ping et al., A highland-adaptation variant near MCUR1 reduces its transcription and attenuates erythrogenesis in Tibetans. Cell Genom 5, 100782 (2025).

32. C. Quan et al., Characterization of structural variation in Tibetans reveals new evidence of high-altitude adaptation and introgression. Genome Biol 22, 159 (2021).

33. G. Ferraretti et al., Archaic introgression contributed to shape the adaptive modulation of angiogenesis and cardiovascular traits in human high-altitude populations from the Himalayas. Elife 12, (2024).

34. W. Zheng et al., Large-scale genome sequencing redefines the genetic footprints of high-altitude adaptation in Tibetans. Genome Biol 24, 73 (2023).

35. F. Bai et al., Ancient genomes revealed the complex human interactions of the ancient western Tibetans. Curr Biol 34, 2594–2605.e2597 (2024).

36. H. Wang et al., Human genetic history on the Tibetan Plateau in the past 5100 years. Sci Adv 9, eadd5582 (2023).

37. S. H. Martin, J. W. Davey, C. D. Jiggins, Evaluating the use of ABBA-BABA statistics to locate introgressed loci. Mol Biol Evol 32, 244–257 (2015).

38. M. A. Harris et al., The Gene Ontology (GO) database and informatics resource. Nucleic Acids Res 32, D258–261 (2004).

39. W. McLaren et al., The Ensembl Variant Effect Predictor. Genome Biol 17, 122 (2016).

40. K. C. Kondapalli, H. Prasad, R. Rao, An inside job: how endosomal Na(+)/H(+) exchangers link to autism and neurological disease. Front Cell Neurosci 8, 172 (2014).

41. S. Fishilevich et al., GeneHancer: genome-wide integration of enhancers and target genes in GeneCards. Database (Oxford) 2017, (2017).

42. P. Rentzsch, D. Witten, G. M. Cooper, J. Shendure, M. Kircher, CADD: predicting the deleteriousness of variants throughout the human genome. Nucleic Acids Res 47, D886–d894 (2019).

43. G. M. Cooper et al., Distribution and intensity of constraint in mammalian genomic sequence. Genome Res 15, 901–913 (2005).

44. F. Hammal, P. de Langen, A. Bergon, F. Lopez, B. Ballester, ReMap 2022: a database of Human, Mouse, Drosophila and Arabidopsis regulatory regions from an integrative analysis of DNA-binding sequencing experiments. Nucleic Acids Res 50, D316–d325 (2022).

45. C. A. Davis et al., The Encyclopedia of DNA elements (ENCODE): data portal update. Nucleic Acids Res 46, D794–d801 (2018).

46. D. Karolchik et al., The UCSC Genome Browser database: 2014 update. Nucleic Acids Res 42, D764–770 (2014).

47. S. S. Rao et al., A 3D map of the human genome at kilobase resolution reveals principles of chromatin looping. Cell 159, 1665–1680 (2014).

48. A. C. Birkeland et al., Correlation of Crtc1/3-Maml2 fusion status, grade and survival in mucoepidermoid carcinoma. Oral Oncol 68, 5–8 (2017).

49. Z. Safarian et al., CCDC82 and neurodevelopment: a novel genetic variant linked to infantile spasms and hypotonia. BMC Med Genomics 18, 133 (2025).

50. W. Q. Yang et al., THUMPD2 catalyzes the N2-methylation of U6 snRNA of the spliceosome catalytic center and regulates pre-mRNA splicing and retinal degeneration. Nucleic Acids Res 52, 3291–3309 (2024).

51. J. C. Keen, H. M. Moore, The Genotype-Tissue Expression (GTEx) Project: Linking Clinical Data with Molecular Analysis to Advance Personalized Medicine. J Pers Med 5, 22–29 (2015).

52. W. Zhang et al., HiLand Resource: A Comprehensive Database of Highland Human Populations. Genomics Proteomics Bioinformatics, (2025).

53. M. Kanehisa, M. Furumichi, M. Tanabe, Y. Sato, K. Morishima, KEGG: new perspectives on genomes, pathways, diseases and drugs. Nucleic Acids Res 45, D353–d361 (2017).

54. L. Qian et al., Peroxisome proliferator-activated receptor gamma coactivator-1 (PGC-1) family in physiological and pathophysiological process and diseases. Signal Transduction and Targeted Therapy 9, 50 (2024).

55. C. Ling et al., Epigenetic regulation of PPARGC1A in human type 2 diabetic islets and effect on insulin secretion. Diabetologia 51, 615–622 (2008).

56. P. Li et al., Metabolomic and microbiome integration of Han-Tibetan and plain-plateau populations. BMC Biol 23, 349 (2025).

57. L. Wang et al., Prevalence and Ethnic Pattern of Diabetes and Prediabetes in China in 2013. Jama 317, 2515–2523 (2017).

58. Y. Li et al., Prevalence of diabetes recorded in mainland China using 2018 diagnostic criteria from the American Diabetes Association: national cross sectional study. Bmj 369, m997 (2020).

59. S. Xu et al., The prevalence of and risk factors for diabetes mellitus and impaired glucose tolerance among Tibetans in China: a cross-sectional study. Oncotarget 8, 112467–112476 (2017).

60. K. Suzuki et al., Identification of 28 new susceptibility loci for type 2 diabetes in the Japanese population. Nat Genet 51, 379–386 (2019).

61. S. A. Liang et al., A refined analysis of Neanderthal-introgressed sequences in modern humans with a complete reference genome. Genome Biol 26, 32 (2025).

62. Y. He et al., Tibetan near-complete pangenome reveals complex variants underlying high-altitude adaptation. bioRxiv [Preprint] (2025). 10.64898/2025.12.16.694547.

63. S. A. Liang et al., Adaptive introgression in Tibetans spans focal and distributed architectures, Data set, Zenodo (2026); 10.5281/zenodo.22763707.

64. L. Li, T. J. Comi, R. F. Bierman, J. M. Akey, Recurrent gene flow between Neanderthals and modern humans over the past 200,000 years. Science 385, eadi1768 (2024).

65. J. Terhorst, J. A. Kamm, Y. S. Song, Robust and scalable inference of population history from hundreds of unphased whole genomes. Nat Genet 49, 303–309 (2017).

66. J. P. Spence, Y. S. Song, Inference and analysis of population-specific fine-scale recombination maps across 26 diverse human populations. Sci Adv 5, eaaw9206 (2019).

67. T. Wu et al., clusterProfiler 4.0: A universal enrichment tool for interpreting omics data. Innovation (Camb) 2, 100141 (2021).

